# Local and global density have distinct and parasite-dependent effects on infection in wild sheep

**DOI:** 10.1101/2024.04.30.591781

**Authors:** Gregory F Albery, Amy R Sweeny, Yolanda Corripio-Miyar, Michael J Evans, Adam Hayward, Josephine M Pemberton, Jill G Pilkington, Daniel H Nussey

**Author notes:** Corresponding authors: Gregory F Albery, 406 Reiss Science Building, 37th and O Streets, NW Washington, DC 20057-1229, 1-202-687-6247; Amy R Sweeny, Alfred Denny Building, Western Bank, Sheffield, UK S10 2TN. Shared lead authorship (Gregory F Albery, Amy R Sweeny). **Author contributions** GFA conceived the study. JMP and JGP led coordination of fieldwork and data collection on St. Kilda. GFA & ARS analysed data with input from DHN. GFA & ARS led writing of the manuscript with input from all authors. **Data accessibility** Data and code for the analyses in this manuscript are available on GitHub at https://github.com/gfalbery/Shensity.

## Abstract

High density should drive greater parasite exposure. However, evidence linking density with infection generally uses density proxies, rather than of individuals per space within a continuous population.

We used a long-term study of wild sheep to link within-population spatiotemporal variation in host density with individual parasite counts. Although four parasites exhibited strong positive relationships with local density, these relationships were mostly restricted to juveniles and faded in adults. Further, one ectoparasite showed strong negative relationships across all age classes. In contrast, population size – a measure of global density – had limited explanatory power, and its effects did not remove those of spatial density, but were complementary. These results indicate that local and global density can exhibit diverse and contrasting effects on infection within populations. Spatial measures of within-population local density may provide substantial additional insight to temporal metrics based on population size, and investigating them more widely could be highly revealing.

## Introduction

An animal’s infection status is driven by its exposure to pathogens, in concert with its susceptibility to infection once exposed [1]. Individuals living in areas of greater population density generally encounter each other more frequently, resulting in a higher *per capita* exposure rate that drives greater prevalence of directly transmitted parasites [2–5]. Additionally, when density drives individuals to share the same space at higher rates, it could likewise drive higher indirect contact rates, leading to greater prevalence of indirectly transmitted parasites; however, density-contact relationships (and therefore density-infection relationships) are likely to differ substantially for parasites of different transmission modes [2]: for example, individuals may be more likely to avoid direct contact while nevertheless sharing space, which could drive more positive density dependence of indirectly than directly transmitted pathogens (Albery et al., in prep). Alternatively, animals may be able to more easily identify areas of environmental parasite transmission than infected conspecifics, which would drive the reverse. Identifying these relationships is important for accurately modelling disease dynamics in many contexts [2,6], and can influence the choice of available interventions. For example, where a disease is transmitted by density-independent interactions, culling the host is unlikely to be effective in reducing its prevalence [7]. Nevertheless, despite a great many studies that employ density dependence functions (reviewed in [2]), there exist relatively few empirical examples of density driving greater infection in individual wild animals. The evidence that exists is often phenomenologically derived or relies on between-species comparisons (e.g. [8–10]), which can be fraught with compensatory evolutionary changes e.g. in social structure [11]. More simply, much of this evidence uses metrics like population size [12,13], or social connectedness metrics like group size [8]. While social contact rates will often correlate positively with density [14–16], social and spatial behaviour can differ inherently and can break this expectation in complex ways, such that using “purely social” metrics may not detect effects of density (i.e., “individuals per space”) *per se*. As such, it is unclear whether population density drives infection within animal systems.

Investigating density itself (i.e., individuals per space, rather than measures of pure sociality) is important because there are a variety of reasons to expect that density will not show a positive relationship with infection [17,18]. For example, habitat selection likely causes individuals to inhabit areas with abundant resources, which creates a positive relationship between nutrition and density; if nutrition results in improved immune resistance [19], this could drive a negative relationship with parasite count – unless the availability of the added resources is cancelled out by greater competition [17]. Reciprocally, competition for resources in a given area is likely to be better approximated by global density, as measured by population size, such that fitting population size as a metric picks up competition for resources more than (or over and above) greater contact rates. These and related processes (see [17,18] for exhaustive lists) could manifest differently for different parasites or host groups within a given population, but such variation in relationships has never been shown and has rarely been investigated. In European badgers (*Meles meles*), social contact metrics showed no relationship with any of five investigated parasites, but a within-population socio-spatial density metric showed a consistent negative linear relationship with four parasites of different transmission modes and host specificities [18]. This negative density dependence was the likely result of parasite avoidance behaviours in space [20,21]. Other studies have shown that density-infection relationships can depend on the timescale at which they are examined [22], as well as other complex patterns of risk [23], few studies have investigated whether density-infection trends diverge across multiple parasites. Understanding why these varied trends might occur is important for understanding how and why parasites regulate spatiotemporal population distributions [17]: if density drives infections, which then influence behaviour and fitness, this process could ultimately feed back to determine who dies where shaping the distribution of the population in space and time.

The wild Soay sheep (*Ovis aries*) of St Kilda have been studied since 1985, with individual-based measurements of behaviour, life history, and parasitism throughout this time [24]. They host a diversity of gastrointestinal parasites, all of which have some environmental phase in between hosts and achieve reinfection through reingestion [25,26]. They also host sheep keds (*Melophagus ovinus*): wingless ectoparasitic flies that achieve transmission through direct contact [27]. Of these parasites, strongyle nematodes are the most important in that they exert strong fitness costs at all life stages [13,28]. Strongyle infection dynamics in the sheep are broadly thought to be driven by population density, where high numbers of grazing sheep produce larger numbers of infectious larvae to be ingested, which then results in greater exposure and therefore greater infection [26]. This mechanism has been linked with greater parasite infection in lambs in high-density years [25,29], and is thought to regulate the population by driving worse condition and greater mortality in these years [26]. Nevertheless, the remaining parasites in the population have yet to be successfully linked with population density [25], and strongyles have not been linked with within-year density metrics or aligned with spatial host distributions; rather, all density analyses have been carried out by linking total population size with infection in a given year, with only one density value applied across the population per year [26]. As such, it remains to be seen how strongyles are driven by density on finer spatiotemporal scales, and whether the density trends are ubiquitous or specific to strongyles. Notably, a previous study found a strong positive correlation between vegetation quality and strongyle count in lambs, which we asserted could be linked to greater density in higher-quality areas [30].

Here, we use 25 years of data in this wild population to examine how a spatial measure of local density drives individual-level infection prevalence and intensity across a range of host age classes, when accounting for global density as approximated by population size. We expected density to be broadly positively correlated with infection because greater host density is associated with greater direct and indirect contact rates, driving greater exposure, and that these effects would be differentiable from – and additional to – the effects of population size.

## Methods

### Study population and parasitology

The Soay sheep are an isolated and unmanaged population that has been monitored since 1985 on the St. Kilda archipelago (57°49′N, 08°34′W, 65 km NW of the Outer Hebrides, Scotland) [24]. Individuals in the Village Bay area of the largest island, Hirta, are individually marked each year and followed longitudinally, with over 95% of individuals in this area being marked at any time [31]. Each spring, lambs are caught shortly after birth in April (typically within a week), marked with unique ear tags and weighed. In August, as many individuals as possible are caught in corral traps over a 2-week period, with 50%–60% of the resident Village Bay sheep population captured each year [24]. Each year, 30 population censuses are carried out by experienced field workers (10 each in spring, summer and autumn); our dataset comprised 961 such censuses. During censuses, fieldworkers follow established routes noting the identity, spatial location (to nearest 100m OS grid square), behaviour and group membership of individual sheep. All sampling was carried out in accordance with UK Home Office regulations under Project Licence PP4825594.

GI parasites were quantified using a modified McMaster technique [26] to enumerate faecal egg counts (FEC, nematodes) or faecal oocyst counts (FOC, protozoans). FEC measures via McMaster techniques have been shown to correlate well to parasite burden in Soay Sheep [26]. FEC/FOC was performed on faecal samples collected in August of each year rectally when animals are captured for morphological measurements and sampling, or from observed defecation within several days of capture where rectal samples could not be obtained. Samples were stored at 4°C until processing which occurred within several weeks of sample collection. Parasite communities of Soay sheep are comprised primarily of gastrointestinal parasites and resemble those of domestic sheep, with the exception of *Haemonchus contortus*. Strongyle nematodes are the dominant group within the Soay parasite fauna, of which there are six known species present in the population, *Teladorsagia circumcincta, Trichostrongylus axei, Tricholostrongylous virtrinus, Chabertia ovina, Bunostomum trignonocephalum*, and *Strongyloides papillosus*. Due to morphological similarities in gastrointestinal nematode eggs which are very difficult or impossible to distinguish, eggs of these species are grouped as a single ‘strongyle’ FEC count within each sample, with the exception of *Strongyloides papillosus*, which is morphologically distinguishable and recorded as present or absent. Soay sheep are also commonly infected with the apicomplexan genus *Eimiera*. Oocysts present in samples from the 11 known species infecting Soay sheep are indistinguishable by eye and grouped as one ‘coccidia’ FOC. Eggs of additional GI helminths *Nematodirus* spp., *Capillaria longipes*, and *Trichuris ovis* are quantified as distinct FECs and the cestode *Moniezia expansa* is scored as present or absent. Although strongyles and coccidia are ubiquitous across age classes and the most common parasite taxa, other taxa are often present at very low prevalence (e.g. *T. ovis & C. longipes*) or only present in specific age categories (e.g. *Nematodirus* spp. in lambs). In addition to gastrointestinal parasites, Soay sheep host the wingless ecotoparasitic fly *Melophagus ovinus* (keds). At each August capture when faecal samples were collected, ked counts were enumerated for each individual via a visual inspection and 1 minute standardised search of the wool on the abdominal region of the animal. FEC/FOC and ked count data used in this study was collected from 1993 to 2017.

### Density measures

We calculated a local density metric for each individual, using all observations of each individual in each year. Our measure of local density followed a method previously described in badgers and deer [14,32], as well as a recent meta-analysis of spatial and social behaviour across wild animal systems (Albery et al., in prep). This approach uses a kernel density estimator with the package `adehabitathr`, taking individuals’ annual centroids and fitting a two-dimensional smoother to the distribution of the data. Individuals are then assigned a local density value (in relative units of individuals per space) based on their annual location on this kernel. This variable therefore represents, for each individual, how many other individuals live in its proximity (i.e., individuals *per space*) in a way that varies predictably across the population and between years.

### Models

Our dataset included 6870 annual measures of 3335 individual sheep, spread across 25 years (1993-2017). We conducted the analysis using R version 4.2.3 [33]. Due to low prevalence, we investigated *Nematodirus* only in lambs and *Capillaria* only in lambs and yearlings (i.e. not in adults).

#### Spatial heterogeneity models

First, to examine spatial patterns of infection, we fitted generalised linear mixed models (GLMMs) using the Integrated Nested Laplace Approximation (INLA) in R [34,35], which is well-suited to identifying spatial autocorrelation and mapping the spatial distribution of wildlife disease [36,37]. We examined each parasite as a count-based response variable with a negative binomial distribution, except the rarer *Capillaria* and *Nematodirus*, for which we used a binomial distribution investigating infection status. We fitted explanatory variables including Sex (two levels: F and M) and Age Category (three levels: Lamb, Yearling, and Adult), with random effects of Individual and Year. To quantify spatial autocorrelation, we fitted a stochastic partial differentiation equation (SPDE) effect, which models samples’ similarity that emerges from their proximity in space. We fitted this effect and compared the fit of the model with and without the effect using deviance information criterion (DIC); a value of -2ΔDIC was taken to denote competitive models – i.e., if adding the SPDE effect reduced DIC by more than 2, it was taken to be significantly spatially autocorrelated. For those parasites with significant spatial heterogeneity, we plotted the distribution of the SPDE effect in space to identify hot- and coldspots of infection.

#### Density models

To identify density-related changes in parasite burden and determine their probable causes, we fitted another selection of GLMMs. These model sets each investigated a different host group: lambs, yearlings, adults, or the population as a whole. We fitted a fixed effect of Sex (two levels: F and M) with random effects of Individual and Year. For the adult models, we included Reproductive Status (Factor with 3 levels: Non-Reproductive Female, Reproductive Female, and Male, instead of Sex); and Age (continuous). For the overall models, we fitted age category (three levels: Lamb, Yearling, and Adult). Finally, we added two measures of density: individual, spatially-defined local density (continuous, standardised to have a mean of 0 and a standard deviation of 1), and population-level measures of global density, defined as the population size in the village bay the following year (continuous, range 211-672, standardised to have a mean of 0 and a standard deviation of 1). Comparing the fit and significance of these two variables would allow us to differentiate effects of local and global density on infection.

## Results

All effect estimates and credibility intervals are given in Supplementary Tables 1-5. Our spatial INLA models found strong spatial autocorrelation in four parasites overall (Figure 1): strongyles (ΔDIC=-6.70); coccidia (ΔDIC=-26.16); *Nematodirus* (ΔDIC=-10.39), and keds (ΔDIC=-34.17). Neither *Strongyloides* nor *Capillaria* were spatially autocorrelated (ΔDIC>-2). Their spatial distributions are displayed in Figure 1. Broadly, there were relatively discordant patterns of the four parasites, but strongyles and *Nematodirus* were generally concentrated in the northeast corner of the population (Figure 1A, C), while coccidia were more concentrated in the southeast (Figure 1B) and keds stood out especially with a strong gradient towards the southwest (Figure 1D).

**Figure 1.**
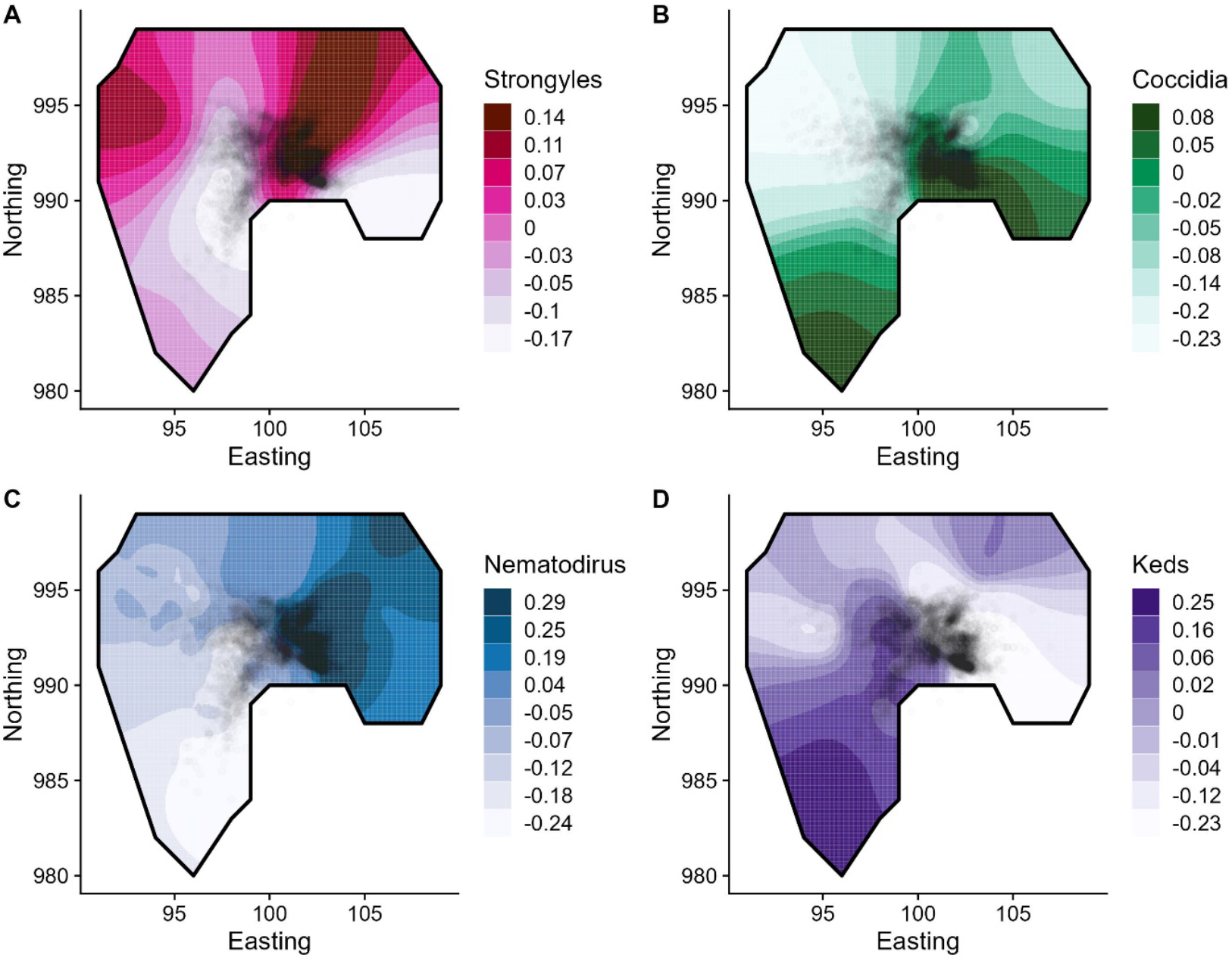
Spatial distributions of four spatially autocorrelated parasites, displayed using the two-dimensional distribution of the SPDE effect from the spatial models including year, sex, age, and random effects of individual and year. Darker colours represent greater parasite count (Panels A, B, D) or prevalence (C), in log-(A, B, D) or logistic (C) units from the mean. Points represent individual average locations based on the population censuses; points are transparent to minimise overplotting. The black line represents the approximate border of the study area.

Fitting individuals’ local density as a fixed effect (overall density distribution displayed in Figure 2A), parasites revealed different relationships with density across age classes (see Figure 2B for all coefficients and P values). 3/6 parasites were positively associated with density in lambs (coccidia, strongyles, and *Nematodirus*), 2/5 in yearlings (*Capillaria* and coccidia), and 1/4 in adults (coccidia). Additionally, counts of keds were negatively correlated with density overall and across all age classes (Figure 2B).

**Figure 2.**
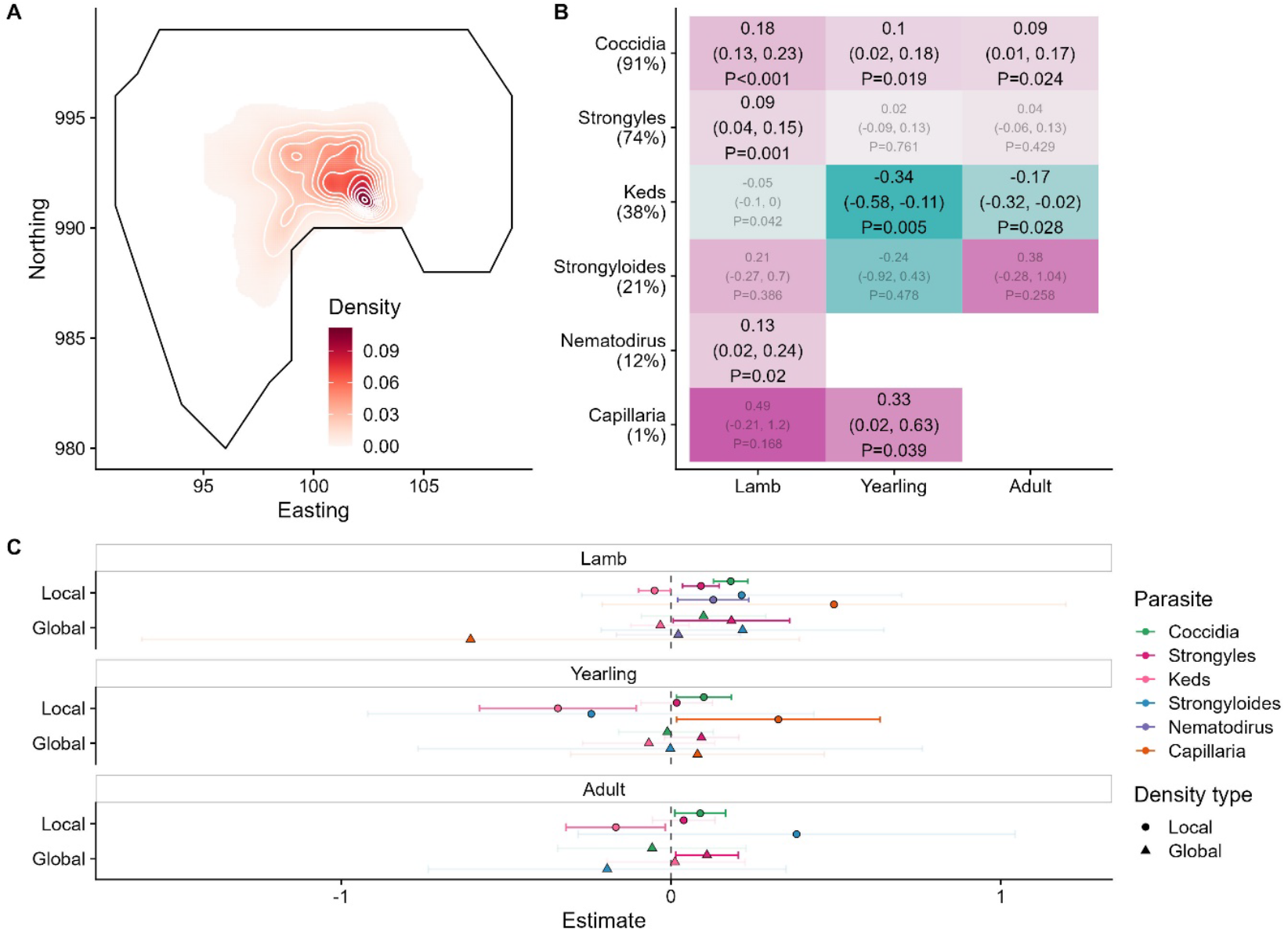
A) Sheep density distribution across the population in space, displayed as an average across the study period. X and Y axes are in easting and northings, in hectares; 1 unit=100m. Darker red colours correspond to greater sheep density in relative units of individuals per space; white contours have been added for clarification. The black line represents the approximate border of the study area, to be aligned with the INLA fields in Figure 1. B) Tile plot depicting the effect sizes for density-infection relationships across parasites and age categories. Tiles are coloured according to the relative positive (pink) or negative (blue) values; a missing tile means the combination was not tested due to low prevalence. Numbers denote the effect size on the log-link scale, with 95% credibility intervals in brackets, and P values. Opaque writing denotes significant effects; transparent writing denotes non-significant effects. C) Effect of global density on parasite count were generally harder to detect than those of local density. Points represent the mean for each effect estimate; error bars denote 95% credibility intervals. All estimates are given on the link scale. Opaque error bars denote estimates that were significant (i.e., their credibility intervals did not overlap with zero); transparent estimates overlap with zero. All effect estimates are given in units of standard deviation.

**Figure 3.**
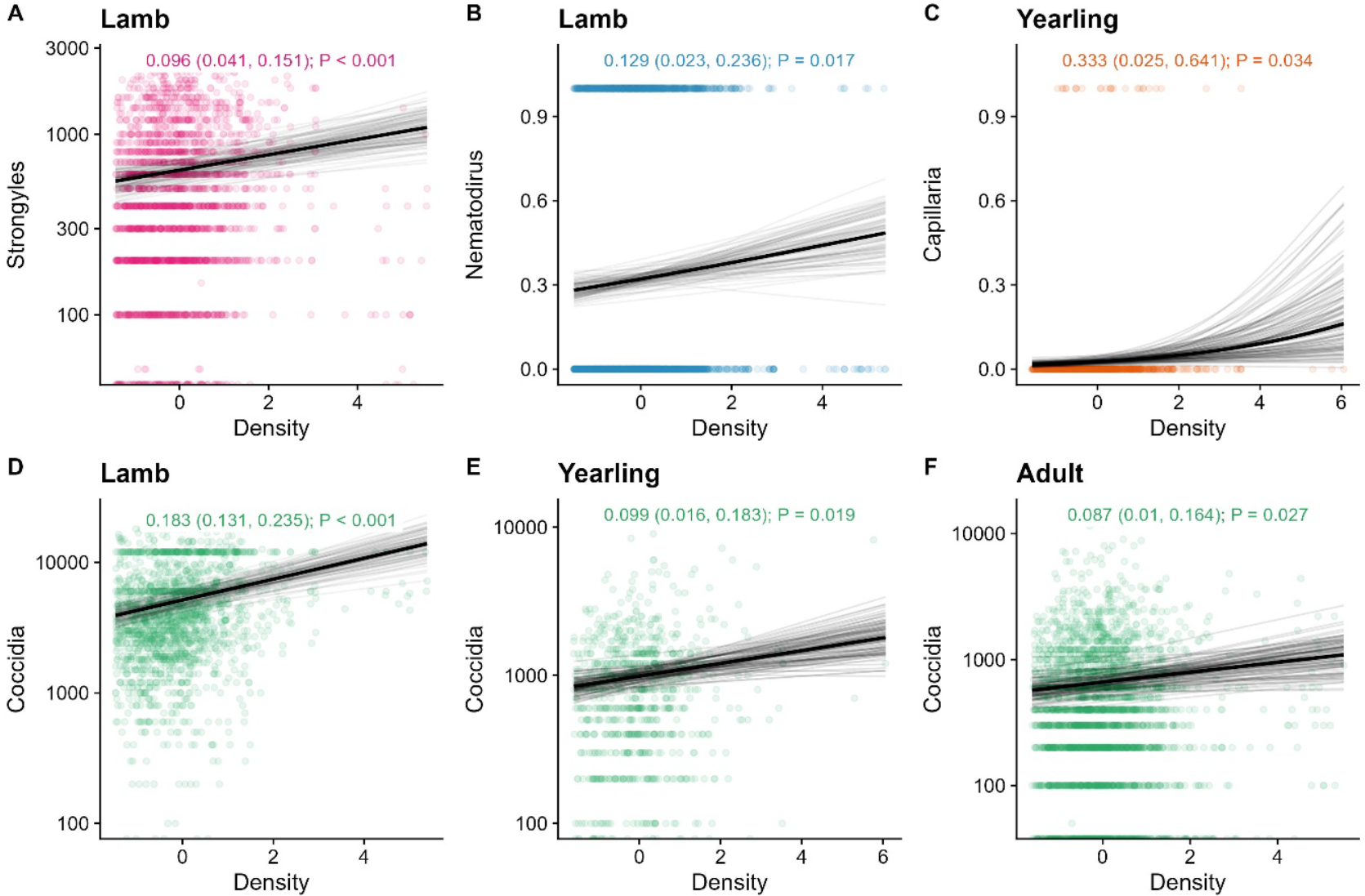
Positive relationships between density and infection in Soay sheep across multiple age categories and parasites. Taken from our density GLMMs, the dark black line represents the mean of the posterior distribution for the age effect estimate; the light grey lines are 100 random draws from the posterior to represent uncertainty. The density effect estimate, credibility intervals, and P values are given at the top of each panel. The points represent individual samples, with transparency to help visualise overplotting. The y axis has been log10-transformed; 0-counts (which are not possible to display on this logged scale) are displayed at the bottom of the graph.

**Figure 4.**
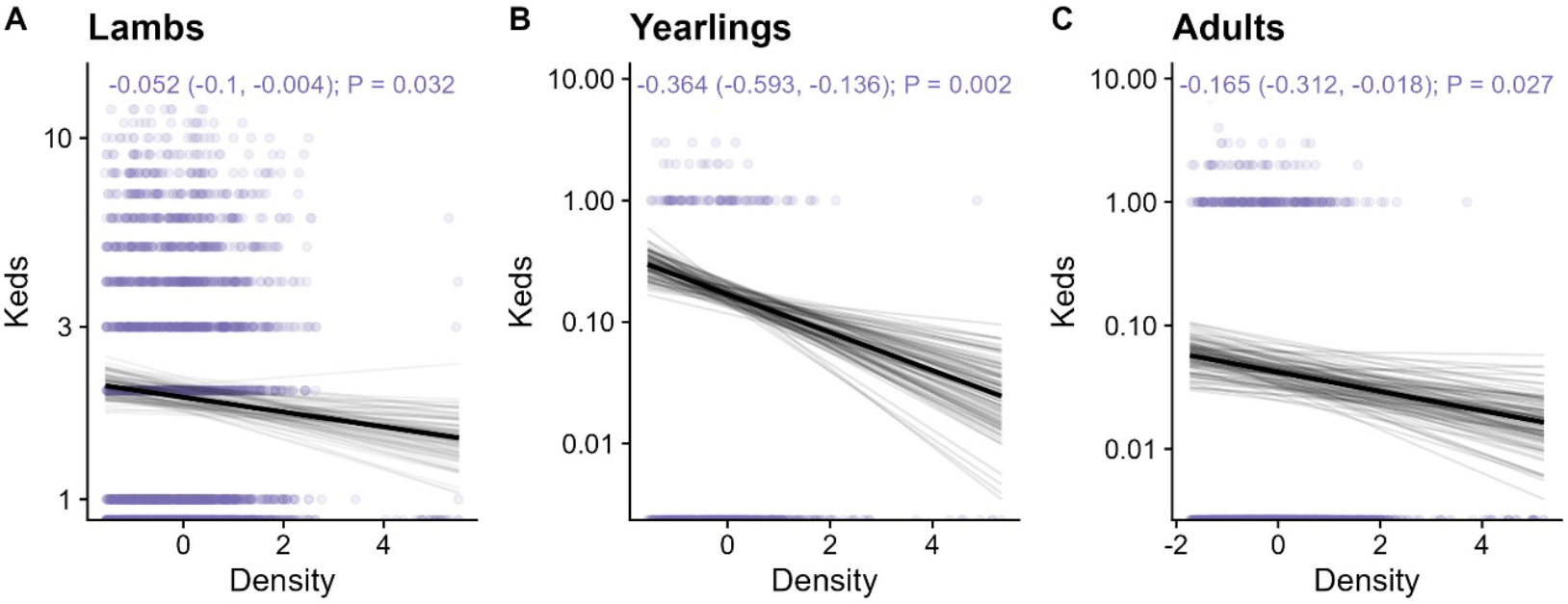
Negative density effects on counts of wingless ectoparasites (Keds, *Melophaga ovinus*) in A) lambs; B) yearlings; and C) adult sheep. Taken from our density GLMMs, the dark black line represents the mean of the posterior distribution for the age effect estimate; the light grey lines are 100 random draws from the posterior to represent uncertainty. The density effect estimate, credibility intervals, and P values are given at the top of each panel. The points represent individual samples, with transparency to help visualise overplotting. The y axis has been log10-transformed; 0-counts (which are not possible to display on this logged scale) are displayed at the bottom of the graph.

In contrast, global density (as approximated by population size the year before sampling) only significantly affected strongyles, in adults (Estimate 0.109; 95% CI 0.014, 0.204; P=0.0241) and lambs (0.183; 0.007, 0.359; P=0.042) but not yearlings (0.092; -0.02, 0.205; P=0.11). Notably, none of these effects removed the effects of local density (see Figure 2C for model estimates); in fact, fitting global density caused the negative effect of local density on ked infection to become significantly negative in lambs. On the contrary, global density’s effect on strongyle infection was additional to local density’s effect in lambs, and provided the only positive density effect on adult infection except for coccidia. Taken together, these results imply that global density and local density are not interchangeable, but complementary sources of information.

## Discussion

As expected, we uncovered strong, largely positive relationships between population density and parasite infection in wild Soay sheep, but with density effects differing substantially across both parasite taxon and host age class. Population density likely drives greater helminth infection by driving greater indirect contact: that is, if more individuals are inhabiting and shedding parasites in higher-density areas, the parasites will achieve a higher concentration on the pasture, thereby driving greater individual-level exposure in these areas. This corroborates prior knowledge concerning the role of population density in driving infection in this population via greater environmental exposure [25,26] – and offers an explanation for the previously observed positive correlation between strongyle count in lambs and preferred vegetation, which is likely to coincide with areas of high host density [30]. However, our use of individual-level (spatial) density metrics allowed us to pick up much finer-scale relationships between density and infection, and thereby detected several more such relationships than previous studies. This demonstrates the value of considering the socio-spatial structuring of a population when examining density-infection interactions (e.g. [18]), rather than focussing solely on variation in population size. Further, while 4/6 parasites showed one or more strong positive relationships with density in lambs or yearlings, there was far weaker evidence of positive density-dependent infection in adults, and ectoparasitic biting fly (“ked”, *Melophaga ovinus*) counts showed unexpected negative correlations with density. Overall, these results serve to demonstrate that socio-spatial measures of population density can influence parasite infection differently across different host and parasite categories within the same population. More generally applying our pipeline across spatially distributed animal populations, as we have previously done in badgers [18], could uncover generalities and contingencies in the factors shaping density-infection relationships [17], allowing us to better understand the mechanisms driving divergent density-infection trends across parasites like these.

Our findings support the role of spatiotemporally heterogeneous population distributions in determining the landscape of infection, and specifically in driving higher burdens in more-social individuals [38]. If parasites have fitness costs, as they are known to in this population [13,26,28], these parasites will likely play an important role regulating the sheep population in space. That is, if density drives higher parasite count and parasites negatively impact fitness, density ultimately drives its own sink factor, determining who dies where. Future studies could attempt to link density and infection with fitness directly to test this spatially explicit population regulation – e.g. via structural equation modelling, which has previously shown that parasites regulate individual reproductive fitness in a similar way in this population [28]. They could further examine whether density-infection relationships depend on resource availability, which should be reduced in higher-density areas and years in this population; this combined exacerbation of exposure and susceptibility will likely exponentially influence parasite count and disproportionately limit the population’s size and density [26,30].

There are several possible explanations for our observation of more positive density-infection relationships in lambs and yearlings than in adults: first, because young individuals have naïve immune systems they are generally highly susceptible to infection, which likely leaves exposure as the main contributor to parasite count [1]. In contrast, between-individual variation in immunity could be complicating relationships between density (i.e. exposure) and infection in older individuals [17,18]. Alternatively, a similar pattern could arise because young individuals are particularly vulnerable to stresses associated with higher density like greater resource competition, reducing their immune resistance and therefore driving greater parasite count [19]. This may be supported by the previously observed positive correlation between strongyle count in lambs and high-quality vegetation, which is likely to coincide with areas of high host density [30]. Alternatively, age-related differences in density-infection patterns could arise through demographic processes linked to those described above: in higher-density areas, heavily infected lambs may die more quickly, leaving behind more resistant lambs and therefore producing a less positive relationship between density and infection at the population level in older individuals (i.e. “selective disappearance” [39]). These findings suggest that density-dependent trends might manifest preferentially for vulnerable (immunologically weaker) classes of hosts, rather than appearing evenly across a population.

Only protozoan coccidia showed consistently positive relationships with density across age categories. Although this could be slightly surprising given that coccidia are generally regarded as more important parasites of juveniles, particularly in ruminants [40], coccidia intensities generally peak across the population in Spring, when lamb density is greatest [41], and independently of the sex and reproductive status effects evident in helminth infections [42]. This observation suggests a combination of lifelong incomplete resistance to infection, such that oocyst concentration on the pasture is a central determinant of individuals’ parasite counts, leading to higher counts in the context of high exposure rates (i.e., in higher density areas). In addition, compared to other parasites, coccidia may be more strongly linked to reduced condition in higher-density areas across all host groups, or their transmission could be more efficient in areas of higher density due to vegetation composition or microclimate, for example. The fitness and condition impacts of coccidia have only been rarely examined in this population [43], but we may be able to interrogate these mechanisms in the future – particularly if combined with analyses of vegetation composition and socio-spatial behaviour [30]. Altogether, the role of coccidia in density dependent regulation of this population could be stronger than previously anticipated.

Contrasting with strongyles and coccidia, we observed consistently negative density effects on counts of ectoparasitic ked flies, which begs explanation, particularly given that direct-contact-transmitted pathogens like keds are generally expected to correlate positively with density [4,8,17]. Because parasite transmission networks are often spatially structured [44], the distribution of environmental factors could influence the way that the sheep interact and transmit keds. For example, sheep may prefer areas with less wind, such that greater wind exposure in lower-density areas drives sheep to more often huddle together or shelter in enclosed spaces. This would lead to higher direct transmission rates *per capita* in low-density areas, producing the observed density-infection relationship. Additionally, at higher densities individuals may paradoxically decrease their direct contact rates or grouping tendencies [45]; along these lines, this divergence in direct and indirect transmission rates with increasing density agrees with our findings elsewhere, which demonstrated stronger indirect than direct transmission is likely to manifest across a range of wild animal systems (Albery et al., in prep).

However, that study showed a saturating positive effect of density on contact in this population specifically, rather than a decrease, so our observation of a negative density-infection trend (rather than a negative one) invites further explanation.

In some scenarios, negative density trends can emerge through encounter-dilution effects, where larger groups subdivide a given burden of mobile parasites – particularly biting insects – such that each individual has fewer parasites [46,47]. This is possible in this case, but given that the keds are not mobile (i.e. without wings) and reside on the sheep themselves, this seems unlikely. Instead, environmental drivers in a given area could drive both lower density and higher ked count. Keds spend much of their lifespan on the skin of the sheep, and they regularly fall off hosts and reside in the environment waiting to encounter another, both of which could leave them relatively exposed to the elements [27]; ked transmission is known to be temperature dependent, supporting this explanation [48]. Sheep may even avoid living in areas that support high ked count due to a desire to avoid infection, driving a negative correlation between density and infection [18,20]. Similarly, ked numbers could be brought down in high-density areas by keds’ natural enemies: keds might be eaten by birds [49], which may result in lower counts in high-density areas if the birds are more likely to inhabit these areas. The keds are also known to transmit (i.e., to host) *Trypanosoma melophagium* blood parasites in this population, and keds elsewhere have been known to transmit bluetongue virus [50] and *Bartonella* bacteria [51]. If these parasites exist at higher prevalence in high-density areas, and have a fitness cost for the vectors, which is fairly common [52,53], they could present a second-degree cost of density for keds; however, highly complex density interactions could arise from this lower intensity of vector infection if they are driven by higher prevalence of their vectored pathogens. Ultimately, future studies of further (internal, rather than ectoparasitic) directly transmitted pathogens, or mapping the distribution of multiple types of between-sheep contacts and their relationships with density, could help to differentiate between these explanations.

Overall, this study provides compelling evidence that density-infection relationships can be strongly contrasting between directly and indirectly transmitted parasites and in hosts of different age classes. Facilitated by more common within-population formulations of such density variables (e.g. [14,18,54]), ecological studies should more often consider that host and parasite biology might introduce strong variation in density-infection relationships, investigating and untangling such contingencies.

## Acknowledgements

ARS was supported by a large NERC grant NE/R016801/1. GFA acknowledges funding from NSF DEB-2211287 and WAI (CBR00730). Field data collection has been supported principally by NERC over many years, with some funding from the Wellcome Trust.

## Supplementary Information

Supplementary Table 1: Estimates and 95% credibility intervals for all response variables and effects for our spatial model set.

Supplementary Table 2: Estimates and 95% credibility intervals for all response variables and effects for our adult model set.

Supplementary Table 3: Estimates and 95% credibility intervals for all response variables and effects for our full-population model set.

Supplementary Table 4: Estimates and 95% credibility intervals for all response variables and effects for our lambs-only model set.

Supplementary Table 5: Estimates and 95% credibility intervals for all response variables and effects for our yearlings-only model set.

